# Two levels of host-specificity in a fig-associated *Caenorhabditis*

**DOI:** 10.1101/261958

**Authors:** Gavin C. Woodruff, Patrick C. Phillips

## Abstract

**Background:** Biotic interactions are ubiquitous and require information from ecology, evolutionary biology, and functional genetics in order to be completely understood. However, study systems that are amenable to investigations across such disparate fields are rare. Figs and fig wasps are a classic system for ecology and evolutionary biology with poor functional genetics; *C. elegans* is a classic system for functional genetics with poor ecology. In order to help bridge these disciplines, here we describe the natural history of a close relative of *C. elegans*, *C.* sp. 34, that is associated with the fig *Ficus septica* and its pollinating *Ceratosolen* wasps.

**Results:** To understand the natural context of fig-associated *Caenorhabditis*, fresh *F. septica* figs from four Okinawan islands were sampled, dissected, and observed under microscopy. *C.* sp. 34 was found in all islands where *F. septica* figs were found. *C.* sp. 34 was routinely found in the fig interior and almost never observed on the outside surface. *Caenorhabditis* was only found in pollinated figs, and *C.* sp. 34 was more likely to be observed in figs with more foundress pollinating wasps. Actively reproducing *C.* sp. 34 dominated younger figs, whereas older figs with emerging wasp progeny harbored *C.* sp. 34 dispersal larvae. Additionally, *C. sp. 34* was observed dismounting from plated *Ceratosolen* pollinating wasps. *C.* sp. 34 was never found on non-pollinating, parasitic *Philotrypesis* wasps. Finally, *C.* sp. 34 was only observed in *F. septica* figs among five Okinawan *Ficus* species sampled.

**Conclusion:** These observations suggest a natural history where *C.* sp. 34 proliferates in young *F. septica* figs and disperses from old figs on *Ceratosolen* pollinating fig wasps. The fig and wasp host specificity of this *Caenorhabditis* is highly divergent from its close relatives and frames hypotheses for future investigations. This natural co-occurrence of the fig/fig wasp and *Caenorhabditis* study systems sets the stage for an integrated research program that can help to explain the evolution of interspecific interactions.

## Background

Interactions at a broad range of scales structure the organization of biological systems. Within ecology, the biotic environment is a major determinant of the distribution and abundance of both species and communities, and so understanding the origins and maintenance of interspecific interactions is a key goal within the field. Yet, interspecific relationships taken as an aggregate are composed of millions of interactions between individual organisms[1, 2], and the nature of those individuals is in turn strongly dependent upon the interactions of thousands of genetic elements comprising their overall genetic composition[3]. Thus a thorough explanation of how and why species interact with one another is ultimately dependent upon information about the genetic bases of such interactions, of which we currently know very little. A full analysis of all of these interactions, from gene to ecosystem, requires the development of study systems in which the power of modern genetic approaches can be used within the context of a compelling ecological circumstance. Here we begin to establish such a system using a newly discovered nematode species that lives in association with the classic fig-fig wasp ecological system[4].

Eukaryotic laboratory model systems have been rightly heralded for their contributions to our understanding of genetics[5-7]. However, only a fraction of their genes are annotated, and there are thousands of genes that as of yet have no known function [8]. Understanding the natural ecological functional context of these genes holds the potential to unlock this mysterious fraction of the genome [8]. Conversely, an understanding of the molecular biology of gene function can be used to inform ecology and evolutionary biology—those interested in the molecular basis of adaptive traits (such as the wing patterns of *Heliconius* butterflies[9] or coat color in crows[10]), physiological systems that structure species distributions[11-13], and the underpinnings of host-microbe interactions[1]) all need functional genetic tools to address their questions [14]. Are such tools also needed to understand the interspecies interactions that underlie most ecological theory?

Successfully traversing these broad fields requires the development of appropriate study systems—particularly systems wherein questions spanning multiple levels of biological organization can be simultaneously addressed. And although there are systems with compelling ecology and evolution (such as *Heliconius* [15], ants/acacias [16], and Darwin’s finches [17]) and systems with well-established and powerful functional genetics (such as fruit flies[6], yeast[5], and worms[7]), systems with a good knowledge of both are rare. The development of good functional genetics in established ecological systems [8] and/or the development of good ecology in established genetic systems[14] is necessary to bridge these gaps.

A classic system for coevolutionary studies is the fig microcosm [4]. The subject of decades of research efforts [18-20], this system has revealed important advances regarding mate competition [21-23], sex ratio allocation [22, 24], and the maintenance of interspecific interactions [25], among others. Furthermore, this system entails a textbook mutualism in figs and their associated wasps: figs need wasps for pollination, and wasps lay their eggs in fig ovules [4]. This system is amenable to experimental manipulation in the field, and evolutionarily-relevant measurements such as the number of seed, wasp progeny, and wasp foundresses are easily ascertained [4]. Thus, this is a powerful system for investigating a number of fundamental questions in ecological and evolution. Likewise, a classic model genetic system is the roundworm *Caenorhabditis elegans.* Like most genetic models, it is easy to rear in the laboratory and is amenable to sophisticated genetic manipulations. Furthermore, the background knowledge concerning its molecular, cellular, and developmental biology is simply vast— we arguably know more about this species than any other metazoan.

Recently, the nematode *Caenorhabditis* sp. 34, a novel sister species to *C. elegans*, has been discovered inside the fresh figs of *Ficus septica* [26]. As multiple reverse genetic techniques are applicable across the genus [27, 28], this species is particularly well-positioned to connect functional genetics with natural ecology. To this end, here we describe the natural ecology context of this fig-associated *Caenorhabditis* through the observation of dissected fresh figs. We examine the extent of *C.* sp. 34 host specificity with both fig and wasp species, the coincidence of worm and fig developmental stages, and the ability of worms to disperse on wasps, with a focus on the implications of these observations for continued studies in both the *C.* sp. 34 and fig/fig-wasp systems.

## Methods

### Collection sites

*C.* sp. 34 was originally isolated from the fresh figs of *Ficus septica* on the island of Ishigaki in Okinawa Prefecture, Japan by Natsumi Kanzaki (Figure 1). To further probe the natural context of this species, *F. septica* figs were sampled from additional Okinawan islands (Figure 2 and Table 1). In May 2015, *F. septica* was sampled from Ishigaki and Iriomote islands, while in n May 2016 sampling of figs was expanded to include the islands of Ishigaki, Iriomote, Miyako, and Yonaguni (Supplemental Table 4). Sampling was also attempted on the islands of Okinawa (main island) and Tarama: *F. septica* was not found at all on Tarama, and although *F. septica* was identified on Okinawa main island, figs were not sampled because no easily-accessible figs could be picked. *F. septica* was typically found at the edge of vegetation on roadsides, but sampling was also performed in the public areas of Banna Park (Ishigaki) and Uenootakejoshi Park (Miyako). In May 2015 and May 2016, additional *Ficus* species were also sampled when accessible figs were found. Images revealing geographic position information of sampled plants were generated with Mapbox.

### Figs dissections and developmental stage classification

Figs were kept refrigerated and dissected <9 days after sampling. Figs were cut into four pieces in tap water in 60 mm petri dishes. In 2015, figs were only scored for *Caenorhadbitis* presence and fig pollination status. In 2016, figs were additionally scored for fig developmental stage, wasp foundress number, and surface nematodes. Unless otherwise noted, the data reported in this study are derived from the larger 2016 set. A fraction of *F. septica* figs (131/250 dissected figs) were initially washed with tap water before dissection in order to interrogate the presence of fig surface nematodes. Dissected figs were then assayed for fig developmental stage, foundress number (in only 169/250 of dissected *F. septica* figs), and *Caenorhabditis* presence under a dissection microscope. *Caenorhabditis* exhibits a stereotypical pharyngeal morphology that was used for genus identification [29]. Figs were binned into five stages based on fig wasp development (inspired by the system developed in [19]; Figure 3a-e): not pollinated (Stage 1), pollinated with no apparent developing wasps (Stage 2), developing wasp progeny apparent (Stage 3), wasp progeny emerging (Stage 4), and post-wasp emergence (Stage 5). In figs where foundress wasps were unambiguous, they were counted. *C.* sp. 34 animals were binned into reproductive phase (L3 stage, L4 stage, and adult; Fig. 1b-c) or dispersal phase (Figure 1d). L1 and L2 stage animals were observed but not noted as they tended to coincide with adult animals and were more difficult to morphologically identify. The dispersing morphotype (Figure 1d) that dominated later stage figs (Fig. 3f) was confirmed to be *Caenorhabditis* in the field via pharynx morphology under compound light microscopy, COI sequencing with phylogenetic analysis (Figure 4), and their development into reproductive stage *Caenorhabditis* under culture conditions (Figure 5). As stress conditions can promote both L1 arrest and dauer larva formation in *Caenorhabditis* [31], and the microscopic power necessary to identify key morphological features of dauer larvae [32] was not available in the field, dispersing animals could not be assigned to such specific developmental stages. This is particularly relevant given the extreme morphological divergence of *C.* sp. 34 [26]. Furthermore, specific *Caenorhabditis* species assignment is problematic as mating tests are typically necessary for species assignment in *Caenorhabditis* [30]. However, it is highly likely that these animals are C. sp. 34 as they share the same specific ecological niche, geographic locality, morphology (Figure 1), and nearly identical COI DNA sequences with the *C.* sp. 34 reference genome (Figure 4). Regardless, reproductive or dispersal *Caenorhabditis* were noted as “abundant” if ≥20 individuals were observed and “rare” if <20 individuals were observed. Dissected figs were observed under a Nikon SMZ-2 dissection microscope, and pharynx morphologies in young larvae were observed with mounted live specimens under a AmScope M100C-LED compound light microscope. DNA sequencing and phylogenetic analysis

**Figure 1.**
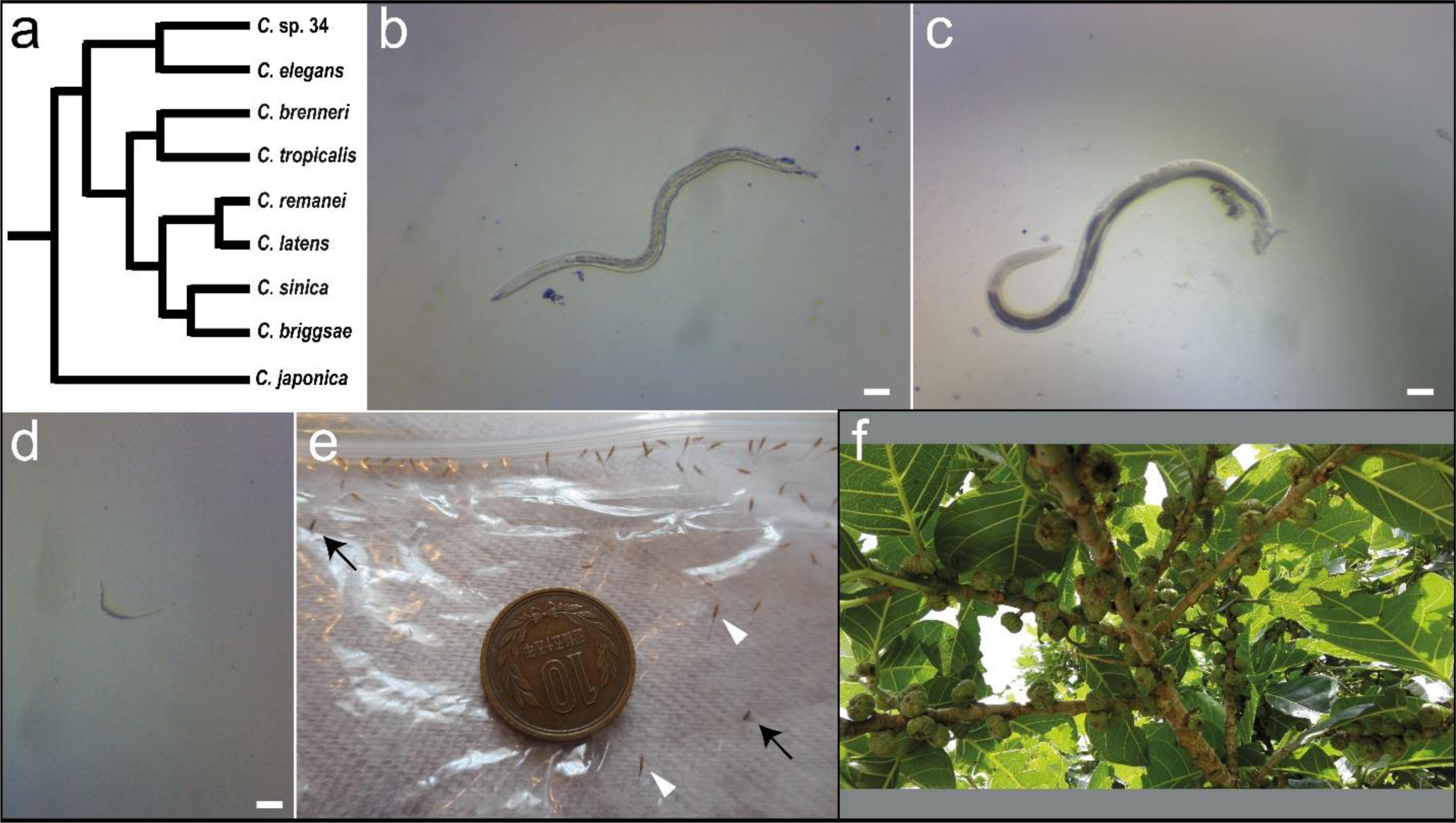
*C.* sp. 34 is associated with fresh *Ficus septica* figs and fig wasps. (a) A cladogram revealing the evolutionarily relationships of *Caenorhabditis*, following [26]. The fig-associated *C.* sp. 34 is among the closest known relatives of the important model organism, *C. elegans.* This reduced figure excludes many known species in this group [30]. (b) An adult *C.* sp. 34 female isolated from a fresh *F. septica* fig. (c) An adult *C.* sp. 34 male isolated from a fresh *F. septica* fig. (d) A dispersal phase *C.* sp. 34isolated from a fresh *F. septica* fig. All scale bars in (b-d) are 100 microns. (e) Fig wasps emerging from fresh *F. septica* figs. Figs were sealed in a plastic bag, and emerging wasps were trapped. Black arrows emphasize female *Ceratosolen* pollinating wasps. White arrowheads highlight *Philotrypesis* parasitic wasps. The ten Japanese yen piece for scale is 23.5 mm in diameter. (f) A *F. septica* plant.

**Figure 2.**
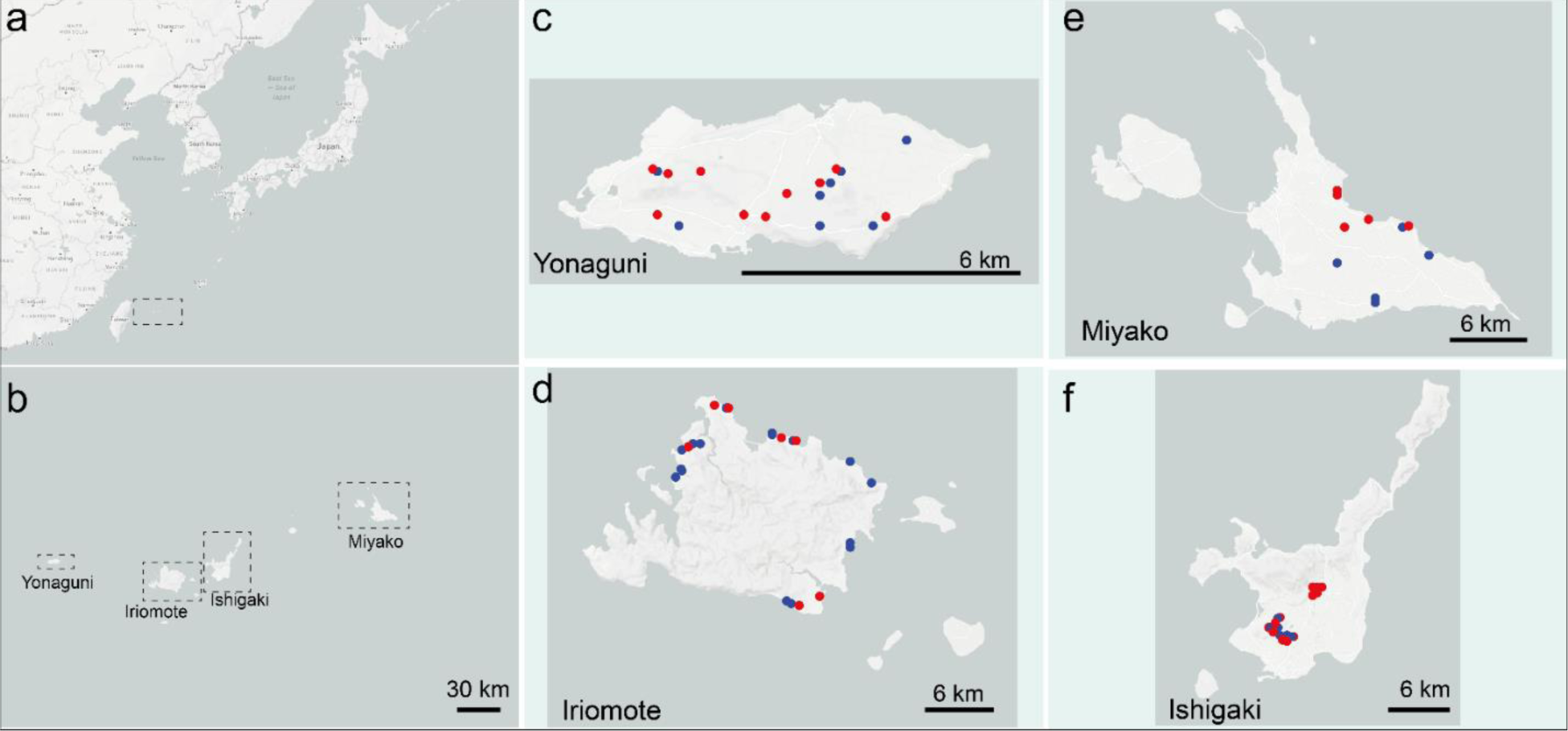
*Ficus septica* fig collection sites in 2016. (a-b) Figs were collected in four of the Sakishima Islands (a, boxed region) of Okinawa Prefecture, Japan: Yonaguni (c), Iriomote (d), Miyako (e), and Ishigaki (f). Blue circles represent positions of *F. septica* plants where *C.* sp. 34 nematodes were found, and red circles denote positions of *F. septica* plants where *C.* sp. 34 nematodes were not found in dissected figs.

*Ficus*, wasp, and nematode species were initially identified via morphological characteristics. Subsequently, DNA was isolated from some ethanol-preserved, *F.* septica-derived wasp and nematode specimens and sequenced to verify genus identity. For wasp samples, preserved animals were washed three times in PBS and subsequently crushed with a pestle in a 1.5 mL Eppendorf tube. DNA was then isolated from the suspension with a Qiagen Blood and Tissue DNeasy kit. For worm DNA samples, preserved single individuals were washed three times in PBS and digested with 5% Proteinase K in Tris-EDTA buffer for 1 hour at 58°C. This solution was immediately used for PCR after a 10 minute, 95°C incubation for enzyme deactivation. For wasp and nematode identification, the mitochondrial cytochrome oxidase I (COI) locus was amplified with primers LCO1490 (5’-GGTCAACAAATCATAAAGATATTGG-3’) and HCO2198 (5’-TAAACTTCAGGGTGACCAAAAAATCA-3’)[33]. PCR reactions were performed with the New England BioLabs Phusion High Fidelity PCR kit. For all reactions this thermocycler program was implemented: 98°C for 10 min. initial denaturation; 98°C for 10 sec. denaturation; 45°C for 30 sec. annealing; 72°C for 30 sec. extension (37 cycles); 72°C for 10 min. final extension. Sanger sequencing was performed by Genewiz. Sequences were then queried with BLAST to the NCBI GenBank database to identify closely related taxa. COI sequences of the fig-associated wasps *Ceratosolen bisculatus* (GenBank accession AF200375), *Apocrypta bakeri* (GenBank accession KF778385), and *Philotrypesis quadrisetosa* (GenBank accession JQ408682), the known fig-associated nematodes *Parasitodiplogaster salicifoliae* (GenBank accession KP015022) and *Schistonchus guangzhouensis* (GenBank accession EU419757), and the marine rhabditid *Litoditus marina* (which was a high BLAST hit for an unidentified nematode species found among our preserved specimens, GenBank accession KR815450). Sequences of the phylogenetically informative taxa *Pristionchus pacificus, C. japonica*, and *C. elegans* were retrieved from WormBase [34]. The *C.* sp. 34 COI sequence was retrieved from the genome assembly (https://www.ncbi.nlm.nih.gov/nuccore?term=382947%5BBioProject%5D). Sequences were aligned with MUSCLE [35], and a maximum likelihood phylogenetic analysis was performed with RaxML[36] under a GTR-gamma model with 1000 bootstrap replicates. Analyses were performed with (335 bp alignment) and without (344 bp alignment) wasp taxa.

**Table 1.** *C.* sp. 34 occupancy in *Ficus septica* figs s in 2016.

|  | Yonaguni | Iriomote | Ishigaki | Miyako | Total |
| --- | --- | --- | --- | --- | --- |
| **Number of plants sampled** | 23 | 26 | 26 | 10 | 84 |
| **Number of plants with *C. sp. 34*** | 10 | 18 | 11 | 5 | 44 |
| **Fraction of plants with *C. sp. 34*** | 0.43 | 0.69 | 0.42 | 0.5 | 0.49 |
| **Number of figs sampled** | 49 | 86 | 37 | 79 | 250 |
| **Number of figs with *C. sp. 34*** | 22 | 49 | 8 | 16 | 95 |
| **Fraction of figs with *C. sp. 34*** | 0.45 | 0.57 | 0.22 | 0.21 | 0.38 |

### Wasp capture, nematode dispersal observations, figs temperature measurements

Parasitic and pollinating fig wasps emerging from intact *F. septica* figs were caught in a plastic bag (Figure 1e). These insects were then killed and placed on Nematode Growth Medium (NGM) agar plates seeded with *E. coli* OP50 bacteria [37]. Plates were monitored for disembarking nematodes three hours and two days after plating. Nematodes of a given morphotype were confirmed to be *C.* sp. 34 via pharyngeal morphology and, in some cases, subsequent development into reproductive phase *C.* sp. 34 (Figure 5).

Additionally, interior and exterior *F. septica* figs temperatures were measured with a DeltaTrack needle thermometer. Each interior measurement was performed on one fresh fig on the tree, and 4-5 figs were measured per plant. These data were taken from about 11:30 AM to 1:30 PM on May 15, 2016 on Yonaguni Island.

**Figure 3.**
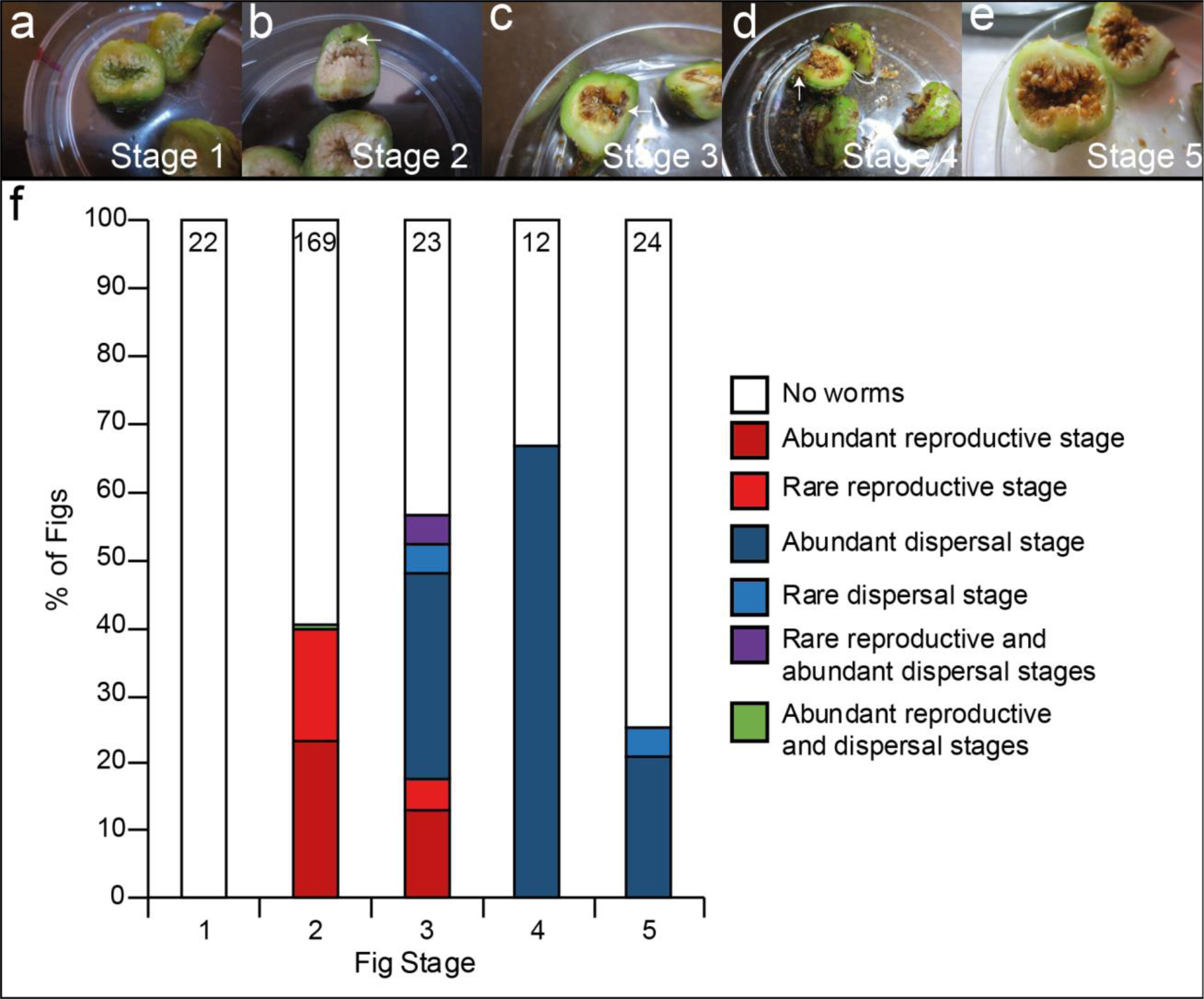
*C.* sp. 34 proliferates in early-stage figs and disperses in late-stage figs. (a-e) Dissected figs were binned into five developmental stages based on wasp presence and developmental progression: (a) not pollinated (Stage 1), (b) pollinated with no apparent developing wasps (Stage 2, arrow noting foundress pollinating wasp), (c) developing wasp progeny apparent (Stage 3), (d) wasp progeny emerging (Stage 4, arrow noting emerging wasp progeny), and (e) post-wasp emergence (Stage 5). The presence of abundant (≥20 individuals) or rare (<20 individuals) reproductive stage or dispersal stage *C.* sp. 34 were noted in each dissected fig (see methods). (f) Frequency of observed *C.* sp. 34 developmental stage by fig developmental stage. Reproductive *C.* sp. 34 predominates in Stage 2 and Stage 3 figs, whereas dispersal *C.* sp. 34 dominates in Stage 4 and Stage 5 figs. *C.* sp. 34 was not observed in figs that were not pollinated. The number of figs dissected per stage is noted at the top of each bar. Adult and dispersal *C.* sp. 34 frequencies were different between fig stages (G-test of independence p-values<0.001 for both adult and dispersal types). Fisher’s Exact Test p-values for all pairwise comparisons can be found in Supplemental Tables 7-8.

**Figure 4.**
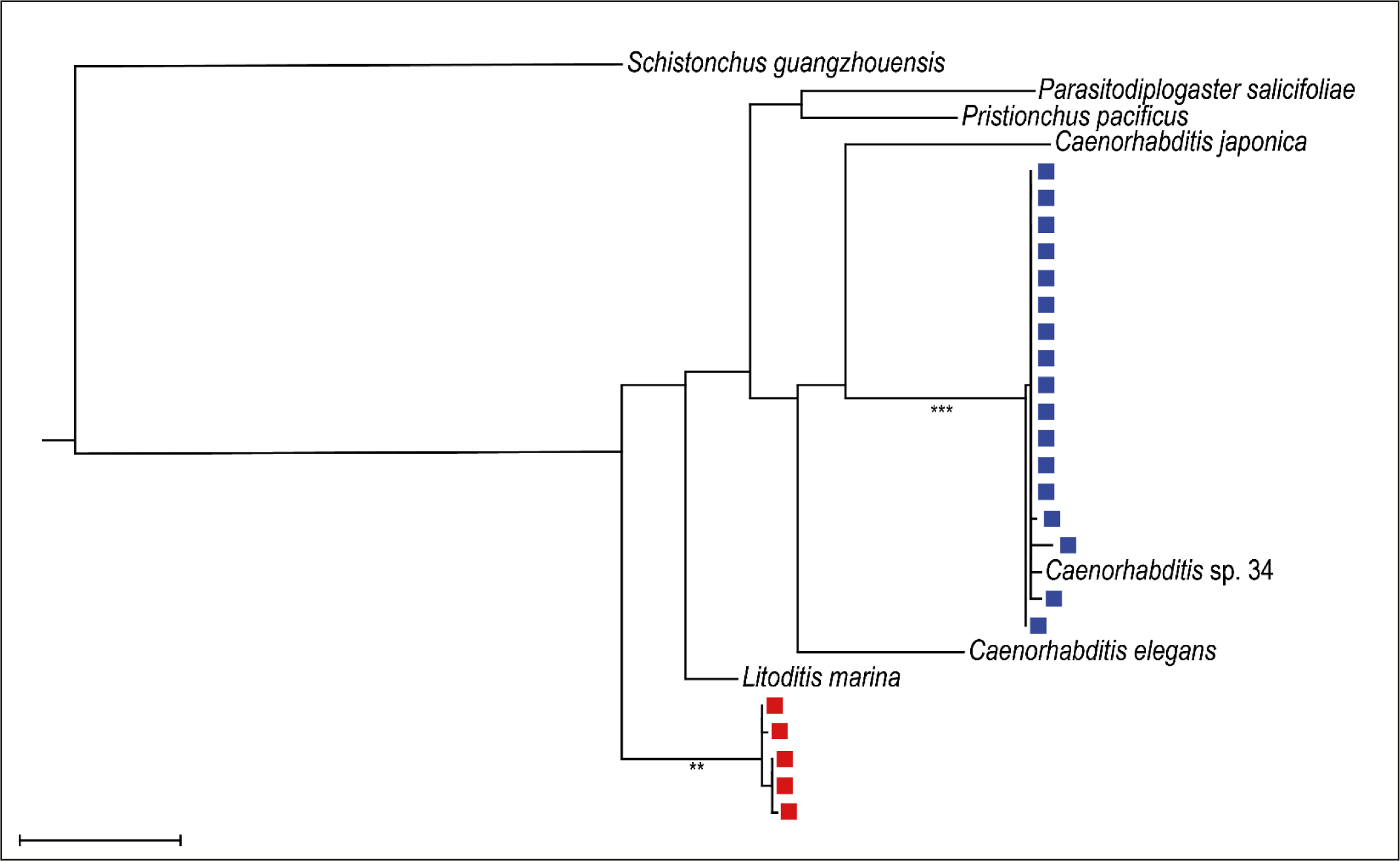
Phylogenetic analysis reveals dispersal larvae share high sequence similarity with *C.* sp. 34. A COI maximum likelihood tree including preserved single dispersal larvae (squares) isolated from a *Ficus septica* fig as well as phylogenetically and ecologically relevant nematode sequences retrieved from GenBank and WormBase (italics). *Parasitodiplogaster* is a known diplogastrid parasite of Ficus-associated wasps [38], and *Schistonchus* is a known tylenchid (clade IV [39]) *Ficus* plant parasite [40]. Although this analysis is insufficient to resolve known *Caenorhabditis* phylogeny [41], 17 individuals (blue squares) share near identical sequence with *C.* sp. 34. An unidentified nematode species (red squares) was also observed; this species is unlikely to be *Schistonchus* or *Parasitodiplogaster* as it does not cluster with representative species in this analysis and returns rhabditid nematodes in a BLAST query to GenBank (top BLAST hits include the marine rhabditid *Litoditis marina* [42], *Phasmarhabditis* sp. [43], and *Acrostichus* sp. [44]; all 88-89% identity). This tree was generated with RaxML under a GTR-gamma model and 1000 bootstrap replicates. Scale bar represents 0.1 substitutions/site. ** = >90% bootstrap support. *** = 100% bootstrap support.

## Results

### *C.* sp. 34 is found inside the fresh, pollinated figs of *Ficus septica*

*C.* sp. 34 was originally isolated from a fresh fig of *Ficus septica* in Okinawa, Japan. To further explore the natural context of this species, *F. septica* figs were collected from additional Okinawan islands (Table 1, Figure 2), dissected, and observed under a dissection microscope for the presence of *C.* sp. 34. *C.* sp. 34 nematodes were found on all four islands where *F. septica* was sampled (Table 1, Figure 2). Although the fraction of *F. septica* plants harboring *C.* sp. 34 in 2016 was largely consistent across islands (G-test of independence p =0.183, Table 1, Supplemental Table 1), the fraction of figs with *C.* sp. 34 showed island-specific differences (G-test of independence p <0.001, Table 1, Supplemental Table 2). Specifically, the *C.* sp. 34 fig occupancy was greater in the two western-most islands of Yonaguni and Iriomote than in the eastern islands of Ishigaki and Miyako (Table 1). These island-specific differences hold even after excluding unpollinated figs (G-test of independence p <0.001, Supplemental Table 3), which were overrepresented on Miyako (Unpollinated fig fraction on Miyako=20/78; unpollinated fig fraction on all other islands=2/172) and were not expected to harbor nematodes (see below). Additionally, few differences were detected between field work seasons (Table 1, Supplemental Tables 4-5). However, *C.* sp. 34 was found less frequently in plants in Ishigaki in 2016 (42% of plants compared to 79% in 2015, Fisher’s exact test p =0.045). Also, between-island differences in fig and plant *Caenorhabditis* occupancy could not be detected in 2015 (Fisher’s exact test p =0.29 and 1, respectively, Supplemental Table 4).

**Figure 5.**
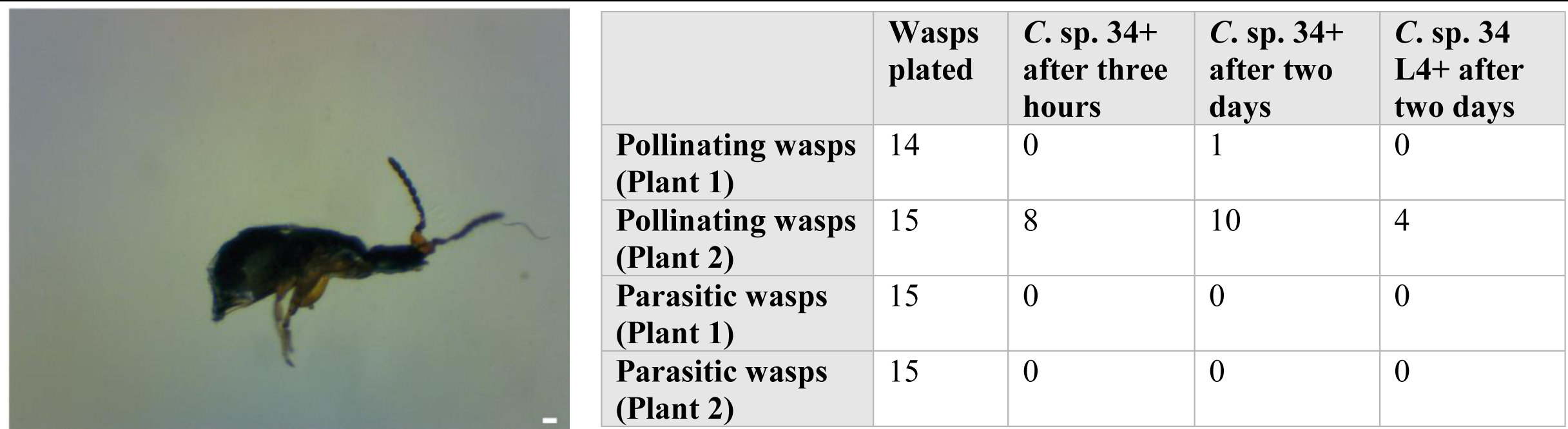
*C.* sp. 34 travels on pollinating fig wasps but not parasitc wasps. Left, a dispersal *Caenorhabditis* nematode dismounts from a pollinating *Ceratosolen* female fig wasp that has been placed on a petri dish. The scale bar represents 10 microns. Right, a table describing wasp carrier data. Fig trees tend to fruit synchronously within a plant but asynchronously between plants [19]. In 2016, two *Ficus septica* plants were observed to harbor figs with actively emerging fig wasps. Emerged fig wasps were caught in a plastic bag, killed, and placed onto agar plates. Plates were subsequently monitored for dismounting *C.* sp. 34 three hours and two days later. Here, numbers represent the number of plated wasps with disembarking *C.* sp. animals. *C.* sp. 34 animals were never seen dismounting from parasitic wasps despite their habitat sharing with pollinating wasps harboring *C.* sp. 34.

*C.* sp. 34 was originally recovered from a dissected fig. To confirm that *C.* sp. 34 proliferates in the interior of the fig and not on its surface, *F. septica* figs were initially washed in tap water and observed under microscopy before and after dissection. The frequency of *C.* sp. 34 observed in washed fresh figs is nearly nonexistent (1 out of 131) compared to that of those subsequently dissected (51 out of 131; Fisher’s Exact test p<0.001). Thus, *C.* sp. 34 is associated with the fig interior and not its surface.

Plants of the genus *Ficus* are renowned for their classic mutualism with pollinating fig wasps [4], and there are a number of Ficus-associated nematodes that require such wasps to complete their life cycle [45]. To interrogate whether this might also hold for fig-associated *Caenorhabditis, F. septica* figs were also queried for their pollination status, which can be ascertained by the presence of developing seed or pollinating wasp progeny. In both field work seasons, *C.* sp. 34 animals were never observed in unpollinated *F. septica* figs (2015: 0/28 unpollinated figs; 2016: 0/22 unpollinated figs). Thus, *C.* sp. 34 likely requires pollinating fig wasps in order to thrive.

In addition to pollination status, the number of foundress pollinating wasps per *F. septica* fig was noted. Typically, female pollinating wasps enter the fig, pollinate it, lay eggs in the fig ovules, and die [4]. In a number of cases, a given fig can have multiple foundresses, which can have profound impacts on wasp population dynamics [23, 24, 46]. Indeed, it was observed that the frequency of *C.* sp. 34 increases with foundress wasp number (Supplemental Figure 1, Supplemental Table 6). The mean foundress number per fig was more than twice as high in figs with *C.* sp. 34 (2.8 wasps, SDM=±1.3, N=72) than in those without (1.1 wasps, SDM=±0.83, N=97; Mann-Whitney U p<0.001). Thus, higher foundress number is associated with *C.* sp. 34 fig occupancy, suggestive that these nematodes disperse on pollinating fig wasps.

### *C.* sp. 34 reproduces in young figs and disperses in old figs

*Caenorhabditis* nematodes can undergo alternative developmental trajectories depending on environmental conditions [47]. If conditions are favorable, animals develop into adults capable of reproduction. But in crowding, starvation, or otherwise stressful conditions, animals develop into the long-lived, stress-resistant dauer larva [47]. It is this dauer stage that is used for dispersal to new food sources in the wild [48]. Previous investigations of fig-associated nematodes have measured the frequency of given nematode developmental stages across fig developmental stages to infer natural histories [45, 49]. To this end, dissected *F. septica* figs were binned into five developmental stages based on wasp presence and development (Figure 3a-e; inspired by the system developed in [19]): not pollinated (Stage 1, Figure 3a); pollinated with no apparent developing wasps (Stage 2, Figure 3b); developing wasp progeny apparent (Stage 3, Figure 3c); wasp progeny emerging (Stage 4, Figure 3d); and post-wasp emergence (Stage 5. Figure 3e). Then, figs were assayed for the presence of rare (<20 individuals) or abundant (≥20 individuals) *C.* sp. 34 reproductive stage (Figure 1b-c) or dispersal stage (Figure 1d) animals. Figure 3f summarizes the results, and it is clear that reproducing *C.* sp. 34 dominate early stage figs. Additionally, dispersal stage *C.* sp. 34 are not found in young figs and rather are only found in older figs that are associated with emerging wasp progeny. Furthermore, subsequent DNA sequencing and phylogenetic analysis using fixed *Ficus-derived* specimens revealed that these dispersal larvae share near identical sequence similarity to sequence retrieved from the *C.* sp. 34 genome assembly (Figure 4), suggestive of identical species status. This distribution of nematode developmental stages then suggests a life cycle wherein fig founders are dispersed by pollinating wasps, proliferate within the young figs, and then generate dispersal forms upon the emergence of wasp progeny.

### *C.* sp. 34 is dispersed by *Ceratosolen* pollinating wasps and not *Philotrypesis* parasitic wasps

To confirm the dispersal of *C.* sp. 34 by fig wasps, emerging *Ceratosolen* pollinating wasps and *Philotrypesis* parasitic wasps were caught in a plastic bag, killed, and placed onto agar plates. Plates were then subsequently monitored at three hours and two days later for the presence of *C.* sp. 34 nematodes. *C.* sp. 34 was observed traveling on pollinating wasps (11/29 wasps; Figure 5) but was never observed on parasitic wasps (0/30 wasps; Figure 5). Of the 11 wasps harboring *C.* sp. 34, there was a median of 2 worms per wasp (range=1-6; Supplemental Figure 2). This was despite both species of wasps emerging from the same figs and the same plant. Thus, *C.* sp. 34 disperses on *Ceratosolen* pollinating fig wasps, and furthermore, *C.* sp. 34 must host-seek within the fig in order to find the proper carrier.

### *Caenorhabditis* has only been found in *F. septica* figs among Okinawan *Ficus*

A number of *Caenorhabditis* species are associated with a variety of plant substrates [50, 51]. However, pollinating fig wasps tend to be associated with only one or two species of *Ficus* [4, 52], which suggests that fig wasp-associated *Caenorhabditis* may also be limited to specific *Ficus* species. To determine if this is so, figs from additional Okinawan *Ficus* species were sampled. Of the nine *Ficus* species reported to be in the sampling locales [53], four species were found with fresh figs aside from *F. septica* (Table 2). No figs aside from *F. septica* were contained *C.* sp. 34 nematodes (Table 2), despite some of these species being known to harbor multiple nematode groups [54, 55]. Thus, this particular fig-associated *C.* sp. 34 is possibly a host specialist and restricted to one species of *Ficus.*

### *F.* septica figs harbor interior temperatures that are comparable to C. sp. 34 lab-rearing temperatures

The environmental parameters defining *Caenorhabditis* ecological niche space are nearly entirely unknown [50]. Among these, temperature influences a multitude of life history traits in *Caenorhabditis*, including survival and reproductive rate [56, 57], as well as the dauer entry switch [58]. To further understand the context of wild *C.* sp. 34, interior *F. septica* live figs and exterior ambient temperatures were measured (Figure 6). Interior fig temperatures (mean=28.7°C, SDM=±1.2, n=39) were on average 2.4°C cooler than exterior temperatures (mean=31.1°C, SDM=±1.5, n=39, t-test p-value<0.001). Interior fig temperatures were comparable to laboratory rearing conditions of *C.* sp. 34, wherein the temperature of 25°C [26] was utilized. Regardless, these observations provide a unique snapshot into the natural context of *C.* sp. 34. Future estimates of additional natural environmental parameters will be essential in informing hypotheses regarding the evolution and ecology of these organisms.

## Discussion

The intricacy of the fig microcosm has facilitated decades of evolutionary and ecological field studies [4, 18]. It harbors a plethora of diverse interspecific interactions: the fig-pollinating wasp mutualism; fig-ant mutualism [59]; fig-nonpollinating wasp parasitism [60]; nematode-wasp parasitism [45]; fig nematode-fig parasitism [40, 61]; and moth-fig parasitism [62]. Figs are also a key resource for over a thousand bird and mammal species, who in turn aid in seed dispersal [63]. As a consequence of this microcosm complexity, this remains an influential and active system for study in ecology and evolution [64-67]. However, none of the species in these communities are particularly amenable to functional genetics and laboratory studies—both of which are crucial for refining the explanatory power of evolutionary science. Conversely, as thousands of genes in multiple long-standing eukaryotic laboratory model systems have no known functions [8], it is likely that their natural ecological contexts (which have often been neglected) will be needed to thoroughly understand their genomes. As a consequence, there have been calls to integrate ecological, evolutionary, and functional genetic approaches[8, 14]. Here, we have described the natural history of C. sp. 34, a close relative of the model genetic organism *C. elegans.* What has been observed in this *Caenorhabditis* study, together with the known biology of the fig microcosm, can then be used to inform hypotheses regarding the evolution of interspecific relationships in both systems.

**Table 2.** *Caenorhabditis* has not been observed in *Ficus* species other than *Ficus septica.*

| **Ficus Species** | **Figs dissected** | **Figs with *C. sp. 34*** | **Figs pollinated** | **Plants Sample** |
| --- | --- | --- | --- | --- |
| **2015** |  |  |  |  |
| *F. superba* | 10 | 0 | -- | 1 |
| *F. microcarpa* | 15 | 0 | 15 | 1 |
| *F. erecta* | 15 | 0 | 3 | 1 |
| **2016** |  |  |  |  |
| *F. variegata* | 10 | 0 | 7 | 2 |
| *F. microcarpa* | 25 | 0 | 25 | 2 |
| *F. erecta* | 36 | 0 | 36 | 2 |

*Caenorhabditis* has not been observed in *Ficus* species other than *Ficus septica*. Non-*F. septica* figs were dissected in May 2015 and May 2016. There have been eight species of *Ficus* aside from *F. septica* reported on these islands [53]. *Caenorhabditis* was not observed in five of these (*F. caulocarpa*, *F. ampelas*, *F. benguetensis*, and *F. virgata* figs were not

*Caenorhabditis* typically proliferates on rotting plants and disperses on invertebrate carriers. And although the features defining niche specialization in this group remain uncertain, it seems clear that there is variation in its extent. Some species appear limited in their geographic range (C. *sinica* has only been found in east Asia [68]), whereas others are globally distributed [50]. Interspecific variation in seasonal predominance of wild populations has been observed, consistent with variation in fitness at different temperatures [51]. Furthermore, different *Caenorhabditis* species have been found associated with different bacterial communities [69], consistent with variation in bacterial preference [70]. There is also interspecific variation in the extent of dispersal carrier specificity. Some *Caenorhabditis* are promiscuous in their choice of carrier; *C. elegans* has been found on snails, slugs, isopods, and myriapods [51]. Other species (such as *C. japonica*, *C. angaria*, and *C. drosophilae*), despite intensive sampling, have only been observed dispersing on one insect species in a highly host-specific manner [50, 71]. The existence of *C.* sp. 34 in the fresh figs of a single species of *Ficus* and observations of its dispersal via pollinating wasps reveals a dramatic shift in substrate from rotting plants to fresh figs. This intimate coupling further reveals an added instance of carrier host-specificity in this group. Further, this niche shift has coincided with extreme morphological and developmental divergence[26], suggesting that this change in natural history has promoted the evolution of novelties within this species. How does the move to the fig microcosm promote such change and otherwise influence their biology?

**Figure 6.**
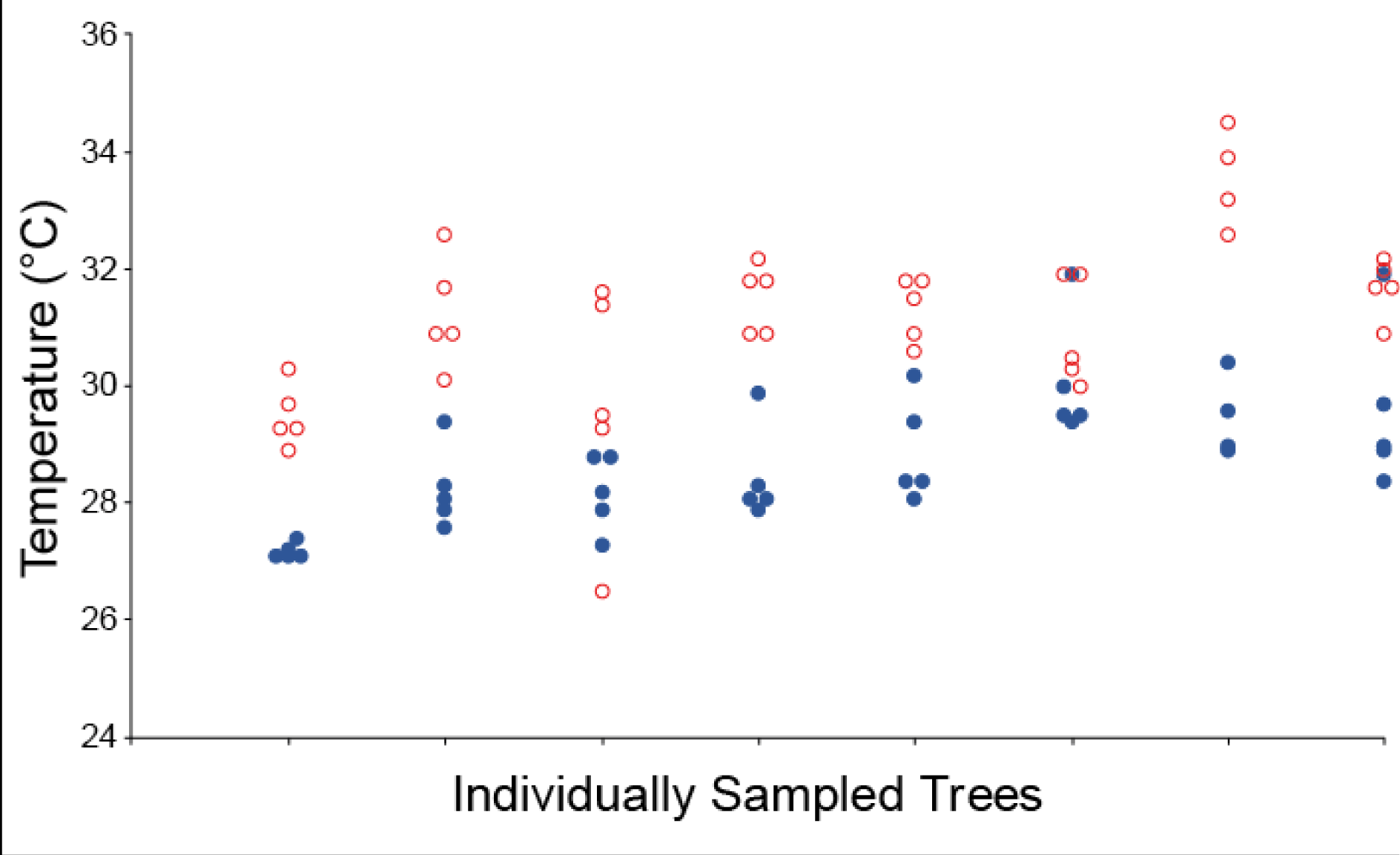
Ambient and interior live *F. septica* figs temperatures. Live *F. septica* figs interiors were measured on eight plants within 1.5 hours in the midday. Open red circles represent exterior temperatures, whereas solid blue circles denote interior fig temperatures. Fig interiors were on average 2.4°C cooler than exterior temperatures (t-test p-value<0.001).

Because *C.* sp. 34 has only been observed dispersing on pollinating fig wasps (Fig. 4), it might be expected that they share similarities in population dynamics. Both pollinating wasp and *C.* sp. 34 founding populations were observed to be quite small (a median of two foundress wasps per fig and two dispersing *C.* sp. 34 per wasp, Supplemental Figures 1-2), consistent with previous observations of inbreeding in pollinating wasps [22, 72]. Variation in founder population size and its inbreeding effects have been shown to have consequences in local mate competition and sex ratio allocation in fig wasps [22, 23]. This may then also hold for *C.* sp. 34, although it is possible that resource availability is different for nematodes (probably bacterial food) and wasps (fig ovules). Furthermore, pollinating fig wasps have been shown to exhibit tremendous dispersal distances [73], which is consistent with low levels of observed population structure for a number of fig wasp populations [72, 74, 75]. These patterns from wasps suggest *C.* sp. 34 may also have unique population genetic features. Male/female *Caenorhabditis* species tend to be incredibly diverse with enormous population sizes, and *C. brenneri* is among the most diverse eukaryotes known [76]. The expected inbreeding in *C.* sp. 34 should reduce diversity, as has been seen in *C. japonica*, another *Caenorhabditis* male/female species with high host-specificity [77]. The selfer *C. elegans* displays reduced diversity, low global population structure, yet high local structure [78, 79]. This is consistent with a boom-and-bust natural history with high migration and largely clonal local populations initiated by single founders [48, 50, 80]. As *C.* sp. 34 is dispersed by wasps that can migrate over long distances while exhibiting small founder populations (Supplemental Figure 1), they may have more population genetic features in common with selfing lineages than expected of a typical gonochoristic *Caenorhabditis* species.

C. sp. 34 also displays differences in developmental timing and developmental decision-making from their close relatives [26]. Their developmental rate is very slow compared to its close relatives, and dauer larvae (an alternative developmental trajectory favored under stress and dispersal conditions) are rarely seen in laboratory populations. Here, we find that reproductive stage animals are enriched in younger figs and smaller, dispersing larvae are found in older figs (Fig. 3). It was not possible to absolutely confirm that these were morphological dauer larvae due to limitations in microscopy in a field setting. However, given that nearly all *Caenorhabditis* observed on invertebrate carriers are in the dauer stage [48], it is likely that animals found in older figs and fig wasps were indeed dauer larvae. Given that figs typically take weeks to develop [18], and that *C.* sp. 34 disperses on pollinating wasps to travel to new figs, it is reasonable to suspect that their divergence in developmental timing and decision-making are related to these features of fig biology. Although it is unclear how many generations are produced within a single fig, C. sp. 34 may have faced selective pressure to slow its developmental rate in order to match progeny production with the timing of wasp emergence. Further, given that dispersal on pollinating wasps is likely critical for *C.* sp. 34 propagation, the decision to enter into dauer may be more dependent on fig and/or wasp chemical cues than those related to stress and population density, which would explain their rarity in laboratory rearing conditions.

The impact of C. sp. 34 on fig and fig wasp fitness remains an open question. Unlike the fig parasite *Schistonchus* [40] and the wasp parasite *Parasitodiplogaster* [38], C. sp. 34 is unlikely to inflict direct harm on figs or wasps as a parasite. This is because *C.* sp. 34 maintains its typical *Caenorhabditis* pharyngeal morphology throughout the reproductive stages observed in fresh figs (plant parasitic nematodes typically have pharyngeal stylets [81]), and proliferative animals are not associated with wasps (Fig. 3; Fig. 5). As a particle feeder, it is possible *C.* sp. 34 eats *Ficus* pollen, thereby affecting host fitness. This seems unlikely, however, as *C. elegans* cannot ingest particles greater than 4 microns in diameter [82], and *Ficus* pollen tends to be larger than this on average [83]. C. sp. 34 may affect pollinator wasp fitness through phoresy by somehow adversely affecting pollinating wasp travel across figs. Considering the size of *C.* sp. 34 dauer larvae (Fig. 1), the low C. sp. 34 dauer larvae load on emerging pollinating wasps (Supplemental Fig. 2), the pervasiveness of phoresy as a dispersal strategy [84], and the contingency of worm success on wasp success in this case, a large cost to wasp dispersal ability also seems unlikely. Instead, *C.* sp. 34 more likely impacts host fitness indirectly through bacteriovory. Microbes harmful or beneficial for fig and wasp fitness could be a major food resource for C. sp. 34. Ants similarly impact fig fitness by discouraging non-pollinating wasps from colonizing figs and are associated with decreased fig herbivory [59]. As measures of fig and wasp fitness (number of seeds and foundress progeny, respectively) are easily obtained [4], and contemporary metagenomic tools can define microbial communities [1], the interplay between C. sp. 34 activity, microbial communities, and host fitness should be able to be interrogated in the future. As our understanding of the *Caenorhabditis-associated* microbiota is rapidly increasing [69, 85, 86], this affords an exciting opportunity for future research.

Notably, *C.* sp. 34 was found dispersing on pollinating *Ceratosolen* wasps, and not *Philotrypesis* parasitoid wasps emerging from figs of the same tree (Figure 5). In contrast to pollinating wasps, who must enter the fig to lay eggs, *Philotrypesis* wasps do not enter the fig and use long ovipositors to lay eggs from the fig exterior [87]. This suggests that dispersing *C.* sp. 34 must discriminate within the fig to find the appropriate carrier. This would likely be a novel behavior, as its close relatives are not fig-associated and tend to be promiscuous in carrier choice [51] (although some preferences in *C. remanei* have been noted [88]). The more distantly-related *C. japonica* has been shown to have behavioral preferences for its shield bug host [89], and similar findings have been shown for *Pristionchus* nematodes and their host beetles [90]. Nematode occupancy biases on pollinating wasps relative to parasitic wasps have been observed in the fig-associated parasitic *Schistonchus* and *Parasitodiplogaster* nematodes [45, 91]. This typical preference for pollinating wasps has been recapitulated in a laboratory framework with *Schistonchus* using traditional chemotaxis assays with wasp-derived volatiles and cuticular hydrocarbons [92]. Similar studies could be extended to the culturable *C.* sp. 34 to interrogate the genetic basis of novel behaviors.

## Conclusion

The elegance of contemporary molecular biology resides in the explanatory power generated by conceptual continuity across multiple hierarchical levels [93] (aka vertical integration [94]). Such continuity is rarely found in evolutionary science—it remains unclear how the disparate pieces of population-level processes, environmental effects, developmental events, and historical contingencies interact to generate diversity in nature. Here, we described the natural history of a close relative of *C. elegans* that is associated with figs and fig wasps. The fig-fig wasp system is a legendary study system in evolution and ecology, and *C. elegans* is a legendary one in model systems genetics. Here then is a serendipitous convergence of research organisms that can facilitate the conceptual connection of their respective disciplines. The functional genetics of *C.* sp. 34 has the potential to inform the molecular basis of how ecologically-relevant phenotypes are generated, whereas the evolution and ecology of the fig system can inform how population-level and environmental forces sort said variation. This all begins with a simple understanding of where and how this organism lives in nature.

## Declarations

### Author contributions

GCW collected and analyzed the data. GCW and PCP wrote the paper.

## Acknowledgements

We thank Natsumi Kanzaki for sharing his expertise in fig-associated nematode field biology. This works was supported by funding from the National Institutes of health to GCW (5F32GM115209-03) and PCP (R01 GM-102511) and from the Japan Society for the Promotion of Science (International Research Fellowship, PE13557) to GCW.

## References

1. Nicholson JK, Holmes E, Kinross J, Burcelin R, Gibson G, Jia W, Pettersson S: Host-gut microbiota metabolic interactions. Science 2012, 336(6086):1262–1267.

2. Thompson JN: The geographic mosaic of coevolution: University of Chicago Press; 2005.

3. Anderson J, Wagner M, Rushworth C, Prasad K, Mitchell-Olds T: The evolution of quantitative traits in complex environments. Heredity 2014, 112(1):4–12.

4. Herre EA, Jandér KC, Machado CA: Evolutionary ecology of figs and their associates: recent progress and outstanding puzzles. Annual Review of Ecology, Evolution, and Systematics 2008:439–458.

5. Botstein D, Fink GR: Yeast: an experimental organism for 21st Century biology. Genetics 2011, 189(3):695–704.

6. Bilder D, Irvine KD: Taking stock of the Drosophila research ecosystem. Genetics 2017, 206(3): 1227–1236.

7. Corsi AK, Wightman B, Chalfie M: A Transparent window into biology: A primer on Caenorhabditis elegans. Genetics 2015, 200(2):387–407.

8. Petersen C, Dirksen P, Schulenburg H: Why we need more ecology for genetic models such as C. elegans. Trends in Genetics 2015, 31(3):120–127.

9. Van Belleghem SM, Rastas P, Papanicolaou A, Martin SH, Arias CF, Supple MA, Hanly JJ, Mallet J, Lewis JJ, Hines HM: Complex modular architecture around a simple toolkit of wing pattern genes. Nature Ecology & Evolution 2017, 1:0052.

10. Poelstra JW, Vijay N, Bossu CM, Lantz H, Ryll B, Müller I, Baglione V, Unneberg P, Wikelski M, Grabherr MG: The genomic landscape underlying phenotypic integrity in the face of gene flow in crows. Science 2014, 344(6190):1410–1414.

11. Storz JF, Runck AM, Sabatino SJ, Kelly JK, Ferrand N, Moriyama H, Weber RE, Fago A: Evolutionary and functional insights into the mechanism underlying high-altitude adaptation of deer mouse hemoglobin. Proceedings of the National Academy of Sciences 2009, 106(34): 14450–14455.

12. Stinchcombe JR, Weinig C, Ungerer M, Olsen KM, Mays C, Halldorsdottir SS, Purugganan MD, Schmitt J: A latitudinal cline in flowering time in Arabidopsis thaliana modulated by the flowering time gene FRIGIDA. Proceedings of the National Academy of Sciences of the United States of America 2004, 101(13):4712–4717.

13. Caicedo AL, Stinchcombe JR, Olsen KM, Schmitt J, Purugganan MD: Epistatic interaction between Arabidopsis FRI and FLC flowering time genes generates a latitudinal cline in a life history trait. Proceedings of the National Academy of Sciences of the United States of America 2004, 101(44): 15670–15675.

14. Bono JM, Olesnicky EC, Matzkin LM: Connecting genotypes, phenotypes and fitness: harnessing the power of CRISPR/Cas9 genome editing. Molecular ecology 2015, 24(15):3810–3822.

15. Jiggins CD: The ecology and evolution of Heliconius butterflies: Oxford University Press; 2016.

16. Heil M, McKey D: Protective ant-plant interactions as model systems in ecological and evolutionary research. Annual Review of Ecology, Evolution, and Systematics 2003:425–453.

17. Grant PR, Grant BR: How and why species multiply: the radiation of Darwin’s finches: Princeton University Press; 2011.

18. Janzen DH: How to be a fig. Annual review of ecology and systematics 1979, 10:13–51.

19. Galil J, Eisikowitch D: On the pollination ecology of Ficus sycomorus in East Africa. Ecology 1968, 49(2):259–269.

20. Weiblen GD: How to be a fig wasp. Annual review of entomology 2002, 47(1):299–330.

21. Moore JC, Loggenberg A, Greeff JM: Kin competition promotes dispersal in a male pollinating fig wasp. Biology letters 2006, 2(1): 17–19.

22. Herre EA: Sex ratio adjustment in fig wasps. Science 1985, 228(4701):896–898.

23. West SA, Murray MG, Machado CA, Griffin AS, Herre EA: Testing Hamilton’s rule with competition between relatives. Nature 2001, 409(6819):510–513.

24. Herre EA: Optimality, plasticity and selective regime in fig wasp sex ratios. Nature 1987, 329(6140):627–629.

25. Jandér KC, Herre EA: Host sanctions and pollinator cheating in the fig tree-fig wasp mutualism. Proceedings of the Royal Society of London B: Biological Sciences 2010, 277(1687): 1481–1488.

26. Woodruff GC, Willis JH, Phillips PC: Dramatic evolution of body length due to postembryonic changes in cell size in a newly discovered close relative of C. elegans. BioRxiv 2017.

27. Nuez I, Félix M-A: Evolution of susceptibility to ingested double-stranded RNAs in Caenorhabditis nematodes. PLoS One 2012, 7(1):e29811.

28. Wei Q, Zhao Y, Guo Y, Stomel J, Stires R, Ellis RE: Co-option of alternate sperm activation programs in the evolution of self-fertile nematodes. Nature communications 2014, 5:5888.

29. Barrière A, Félix M-A: Isolation of C. elegans and related nematodes. WormBook 2014.

30. Félix M-A, Braendle C, Cutter AD: A streamlined system for species diagnosis in Caenorhabditis (Nematoda: Rhabditidae) with name designations for 15 distinct biological species. PLoS One 2014, 9(4):e94723.

31. Baugh LR: To grow or not to grow: nutritional control of development during Caenorhabditis elegans L1 arrest. Genetics 2013, 194(3):539–555.

32. Hu PJ: Dauer. In: WormBook. Edited by Community TCeR: WormBook.

33. Hebert PD, Cywinska A, Ball SL: Biological identifications through DNA barcodes. Proceedings of the Royal Society of London B: Biological Sciences 2003, 270(1512):313–321.

34. Howe KL, Bolt BJ, Cain S, Chan J, Chen WJ, Davis P, Done J, Down T, Gao S, Grove C: WormBase 2016: expanding to enable helminth genomic research. Nucleic acids research 2015:gkv1217.

35. Edgar RC: MUSCLE: multiple sequence alignment with high accuracy and high throughput. Nucleic acids research 2004, 32(5): 1792–1797.

36. Stamatakis A: RAxML version 8: a tool for phylogenetic analysis and post-analysis of large phylogenies. Bioinformatics 2014, 30(9):1312–1313.

37. Brenner S: The genetics of Caenorhabditis elegans. Genetics 1974, 77(1):71–94.

38. Poinar GO, Herre EA: Speciation and adaptive radiation in the fig wasp nematode Parasitodiplogaster (Diplogasteridae: Rhabditida) in Panama. Revue de Nématologie 1991, 14(3):361–374.

39. Blaxter ML, De Ley P, Garey JR, Liu LX, Scheldeman P, Vierstraete A, Vanfleteren JR, Mackey LY, Dorris M, Frisse LM: A molecular evolutionary framework for the phylum Nematoda. Nature 1998, 392(6671):71–75.

40. Davies KA, Ye W, Kanzaki N, Bartholomaeus F, Zeng Y, Giblin-Davis RM: A review of the taxonomy, phylogeny, distribution and co-evolution of SchistonchusCobb, 1927 with proposal of Ficophagusn. gen. and Martinineman. gen.(Nematoda: Aphelenchoididae). Nematology 2015, 17(7):761–829.

41. Kiontke KC, Félix M-A, Ailion M, Rockman MV, Braendle C, Pénigault J-B, Fitch DH: A phylogeny and molecular barcodes for Caenorhabditis, with numerous new species from rotting fruits. BMC Evolutionary Biology 2011, 11(1):339.

42. Derycke S, De Meester N, Rigaux A, Creer S, Bik H, Thomas W, Moens T: Coexisting cryptic species of the Litoditis marina complex (Nematoda) show differential resource use and have distinct microbiomes with high intraspecific variability. Molecular ecology 2016.

43. Ross JL, Malan AP: Nematodes Associated with Terrestrial Slugs. In: Nematology in South Africa: A View from the 21st Century. Springer; 2017: 481–493.

44. Mayer WE, Herrmann M, Sommer RJ: Molecular phylogeny of beetle associated diplogastrid nematodes suggests host switching rather than nematode-beetle coevolution. BMC Evolutionary Biology 2009, 9(1):212.

45. Giblin-Davis RM, Center BJ, Nadel H, Frank JH, Ramírez W: Nematodes associated with fig wasps, Pegoscapus spp.(Agaonidae), and syconia of native Floridian figs (Ficus spp.). Journal of Nematology 1995, 27(1): 1.

46. Herre EA: Population structure and the evolution of virulence in nematode parasites of fig wasps. SCIENCE-NEW YORK THEN WASHINGTON 1993, 259:1442–1442.

47. Fielenbach N, Antebi A: C. elegans dauer formation and the molecular basis of plasticity. Genes & development 2008, 22(16):2149–2165.

48. Félix M-A, Braendle C: The natural history of Caenorhabditis elegans. Current Biology 2010, 20(22):R965–R969.

49. Susoy V, Herrmann M, Kanzaki N, Kruger M, Nguyen CN, Rödelsperger C, Röseler W, Weiler C, Giblin-Davis RM, Ragsdale EJ: Large-scale diversification without genetic isolation in nematode symbionts of figs. Science Advances 2016, 2(1):e1501031.

50. Cutter AD: Caenorhabditis evolution in the wild. BioEssays 2015, 37(9):983–995.

51. Félix M-A, Duveau F: Population dynamics and habitat sharing of natural populations of Caenorhabditis elegans and C. briggsae. BMC biology 2012, 10(1): 1.

52. Molbo D, Machado CA, Sevenster JG, Keller L, Herre EA: Cryptic species of fig-pollinating wasps: implications for the evolution of the fig-wasp mutualism, sex allocation, and precision of adaptation. Proceedings of the National Academy of Sciences 2003, 100(10):5867–5872.

53. Yokoyama J, Iwatsuki K: A faunal survey of fig-wasps (Chalcidoidea: Hymenoptera) distributed in Japan and their associations with figs (Ficus: Moraceae). Entomological science 1998, 1(1):37–46.

54. Kanzaki N, Woodruff GC, Tanaka R: Teratodiplogaster variegatae n. sp.(Nematoda: Diplogastridae) isolated from the syconia of Ficus variegata Blume on Ishigaki Island, Okinawa, Japan. Nematology 2014, 16(10): 1153–1166.

55. Zeng Y, Ye W, Giblin-Davis RM, Li C, Zhang S, Du Z: Description of Schistonchus microcarpus n. sp.(Nematoda: Aphelenchoididae), an associate of Ficus microcarpa in China. Nematology 2011, 13(2):221–233.

56. Byerly L, Cassada R, Russell R: The life cycle of the nematode Caenorhabditis elegans: I. Wild-type growth and reproduction. Developmental biology 1976, 51(1):23–33.

57. Anderson JL, Albergotti L, Ellebracht B, Huey RB, Phillips PC: Does thermoregulatory behavior maximize reproductive fitness of natural isolates of Caenorhabditis elegans? BMC evolutionary biology 2011, 11(1): 157.

58. Golden JW, Riddle DL: The Caenorhabditis elegans dauer larva: developmental effects of pheromone, food, and temperature. Developmental biology 1984, 102(2):368–378.

59. Jander KC: Indirect mutualism: ants protect fig seeds and pollen dispersers from parasites. Ecological Entomology 2015, 40(5):500–510.

60. Borges RM: How to be a fig wasp parasite on the fig-fig wasp mutualism. Current Opinion in Insect Science 2015, 8:34–40.

61. Kanzaki N, Tanaka R, Giblin-Davis RM, Davies KA: New plant-parasitic nematode from the mostly mycophagous genus Bursaphelenchus discovered inside figs in Japan. PloS one 2014, 9(6):e99241.

62. Sugiura S, Yamazaki K: Moths boring into Ficus syconia on Iriomote Island, south-western Japan. Entomological Science 2004, 7(2): 113–118.

63. Shanahan M, So S, Compton SG, Corlett R: Fig-eating by vertebrate frugivores: a global review. Biological Reviews of the Cambridge Philosophical Society 2001, 76(04):529–572.

64. Kjellberg F, Proffit M: Tracking the elusive history of diversification in plant-herbivorous insect-parasitoid food webs: insights from figs and fig wasps. Molecular ecology 2016, 25(4):843–845.

65. Sutton TL, Riegler M, Cook JM: One step ahead: a parasitoid disperses farther and forms a wider geographic population than its fig wasp host. Molecular ecology 2016, 25(4):882–894.

66. Jandér K, Dafoe A, Herre E: Fitness reduction for uncooperative fig wasps through reduced offspring size: A third component of host sanctions. Ecology 2016.

67. Sun B-F, Li Y-X, Jia L-Y, Niu L-H, Murphy RW, Zhang P, He S, Huang D-W: Regulation of transcription factors on sexual dimorphism of fig wasps. Scientific reports 2015, 5.

68. Huang R-E, Ren X, Qiu Y, Zhao Z: Description of Caenorhabditis sinica sp. n.(Nematoda: Rhabditidae), a nematode species used in comparative biology for C. elegans. PloS one 2014, 9(11):e110957.

69. Dirksen P, Marsh SA, Braker I, Heitland N, Wagner S, Nakad R, Mader S, Petersen C, Kowallik V, Rosenstiel P: The native microbiome of the nematode Caenorhabditis elegans: gateway to a new host-microbiome model. BMC biology 2016, 14(1): 1.

70. Glater EE, Rockman MV, Bargmann CI: Multigenic natural variation underlies Caenorhabditis elegans olfactory preference for the bacterial pathogen Serratia marcescens. G3: Genes| Genomes| Genetics 2014, 4(2):265–276.

71. Kiontke K, Sudhaus W: Ecology of Caenorhabditis species. In: WormBook. Edited by Community TCeR: WormBook; 2006.

72. Molbo D, Machado CA, Herre EA, Keller L: Inbreeding and population structure in two pairs of cryptic fig wasp species. Molecular Ecology 2004, 13(6): 1613–1623.

73. Ahmed S, Compton SG, Butlin RK, Gilmartin PM: Wind-borne insects mediate directional pollen transfer between desert fig trees 160 kilometers apart. Proceedings of the National Academy of Sciences 2009, 106(48):20342–20347.

74. Lin RC, YEUNG CKL, LI SH: Drastic post-LGM expansion and lack of historical genetic structure of a subtropical fig-pollinating wasp (Ceratosolen sp. 1) of Ficus septica in Taiwan. Molecular Ecology 2008, 17(23):5008–5022.

75. Kobmoo N, HOSSAERT-MCKEY M, Rasplus J, Kjellberg F: Ficus racemosa is pollinated by a single population of a single agaonid wasp species in continental South-East Asia. Molecular Ecology 2010, 19(13):27001–2712.

76. Dey A, Chan CK, Thomas CG, Cutter AD: Molecular hyperdiversity defines populations of the nematode Caenorhabditis brenneri. Proceedings of the National Academy of Sciences 2013, 110(27): 11056–11060.

77. Li S, Jovelin R, Yoshiga T, Tanaka R, Cutter AD: Specialist versus generalist life histories and nucleotide diversity in Caenorhabditis nematodes. Proceedings of the Royal Society of London B: Biological Sciences 2014, 281(1777):20132858.

78. Barrière A, Félix M-A: High local genetic diversity and low outcrossing rate in Caenorhabditis elegans natural populations. Current Biology 2005, 15(13): 1176–1184.

79. Andersen EC, Gerke JP, Shapiro JA, Crissman JR, Ghosh R, Bloom JS, Félix M-A, Kruglyak L: Chromosome-scale selective sweeps shape Caenorhabditis elegans genomic diversity. Nature genetics 2012, 44(3):285–290.

80. Frézal L, Félix M-A: C. elegans outside the Petri dish. Elife 2015, 4:e05849.

81. Bird DM, Jones JT, Opperman CH, Kikuchi T, Danchin EG: Signatures of adaptation to plant parasitism in nematode genomes. Parasitology 2015, 142(S1):S71–S84.

82. Fang-Yen C, Avery L, Samuel AD: Two size-selective mechanisms specifically trap bacteria-sized food particles in Caenorhabditis elegans. Proceedings of the National Academy of Sciences 2009, 106(47):20093–20096.

83. Wang G, Chen J, Li Z-B, Zhang F-P, Yang D-R: Has pollination mode shaped the evolution of ficus pollen? PloS one 2014, 9(1):e86231.

84. Schulenburg H, Félix M-A: The Natural Biotic Environment of Caenorhabditis elegans. Genetics 2017, 206(1):55–86.

85. Berg M, Stenuit B, Ho J, Wang A, Parke C, Knight M, Alvarez-Cohen L, Shapira M: Assembly of the Caenorhabditis elegans gut microbiota from diverse soil microbial environments. The ISME journal 2016, 10(8):1998–2009.

86. Samuel BS, Rowedder H, Braendle C, Félix M-A, Ruvkun G: Caenorhabditis elegans responses to bacteria from its natural habitats. Proceedings of the National Academy of Sciences 2016, 113(27):E3941–E3949.

87. Cook J, Bean D: Cryptic male dimorphism and fighting in a fig wasp. Animal Behaviour 2006, 71(5): 1095–1101.

88. Baird SE: Natural and experimental associations of Caenorhabditis remanei with Trachelipus rathkii and other terrestrial isopods. Nematology 1999, 1(5):471–475.

89. Okumura E, Tanaka R, Yoshiga T: Species-specific recognition of the carrier insect by dauer larvae of the nematode Caenorhabditis japonica. Journal of Experimental Biology 2013, 216(4):568–572.

90. Hong RL: Pristionchus pacificus olfaction. In: Pristionchus pacificus. Brill; 2015: 331–352.

91. Vovlas N, Larizza A: Relationship of Schistonchus caprifici (Aphelenchoididae) with fig inflorescences, the fig pollinator Blastophaga psenes, and its cleptoparasite Philotrypesis caricae. Fundamental and Applied Nematology 1996, 19(5):443–448.

92. Krishnan A, Muralidharan S, Sharma L, Borges RM: A hitchhiker’s guide to a crowded syconium: how do fig nematodes find the right ride? Functional Ecology 2010, 24(4):741–749.

93. Wagner GP: The current state and the future of developmental evolution. From embryology to evo-devo: A history of developmental evolution 2007:525–545.

94. Lee YW, Gould BA, Stinchcombe JR: Identifying the genes underlying quantitative traits: a rationale for the QTN programme. AoB Plants 2014, 6:plu004.

